# Discrete and Continuous Cell Identities of the Adult Murine Striatum

**DOI:** 10.1101/591396

**Authors:** Geoffrey Stanley, Ozgun Gokce, Robert C. Malenka, Thomas C. Südhof, Stephen R. Quake

## Abstract

The striatum is a large brain region containing two major cell types: D1 (dopamine receptor 1) and D2 (dopamine receptor 2) expressing spiny projection neurons (SPNs). We generated a cell type atlas of the adult murine striatum using single-cell RNA-seq of SPNs combined with quantitative RNA *in situ* hybridization (ISH). We developed a novel computational pipeline that distinguishes discrete versus continuous cell identities in scRNA-seq data, and used it to show that SPNs in the striatum can be classified into four discrete types that reside in discrete anatomical clusters or are spatially intermingled. Within each discrete type, we find multiple independent axes of continuous cell identity that map to spatial gradients and whose genes are conserved between discrete types. These gradients correlate well to previously-mapped gradients of connectivity. Using these insights, we discovered multiple novel spatially localized region of the striatum, one of which contains patch-D2 SPNs that express *Tac1, Htr7*, and *Th*. Intriguingly, we found one subtype that strongly co-expresses both D1 and D2 dopamine receptors, and uniquely expresses a rare D2 receptor splice variant. These results collectively suggest an organizational principal of neuron identity in which major neuron types can be separated into discrete classes with little overlap and no implied spatial relationship. However these discrete classes are then continuously subdivided by multiple spatial gradients of expression defining anatomical location via a combinatorial mechanism. Finally, they suggest that neuronal circuitry has a substructure at far higher resolution than is typically interrogated which is defined by the precise identity and location of a neuron.

## Introduction

Neuronal computation is typically thought of as the interactions of defined circuits between large groups of cells called neuron types, which are interrogated by using marker genes that are expressed only in a given type in a brain region. Single-cell transcriptomics has advanced these efforts by enabling the identification of markers for hundreds of candidate neuron types across many brain regions^1–10^. Broad neuron classes generally form clear, non-overlapping clusters, often distinguishable by a single marker gene^1,11,12^. Within those classes, however, neuron types often overlap or form gradients^1,11,12^ suggesting the existence of a further layer of identity structure within each discrete type^14^. It is not clear how best to analyze single cell data to reveal these differences, or whether this continuous heterogeneity can even be used to classify neurons^13,14^. These controversies are partly due to the possibility that continuous heterogeneities represent transient cell states or genetic differences between animals^13^. More generally, this phenomenon calls into question the paradigms of assigning cell types into discrete identities and defining neuronal circuits by large, homogeneous groups of cells.

We created a single-cell sequencing dataset of a brain region paired with an *in situ* analysis of every subtype that provides a ground-truth reference for which neuronal identities are discrete and which are varying continuously. To analyze this data we developed a novel algorithm that distinguishes between continuous and discrete variation. We used this algorithm to show that striatal heterogeneity is comprised of four discrete subtypes of SPNs; three of these discrete subtypes vary continuously along multiple shared axes of gene expression, and these shared continuous axes encode for spatial location. We also find spatial compartments that are limited to a single discrete type, and we discovered a novel region of the striatum, which we call the ventromedial patch and is composed of D2 SPNs that co-express *Tac1, Penk, Htr7*, and *Th*. We used full-length cDNA to show that a subtype, which we previously referred to as D1-*Pcdh8*, co-expresses *Drd1a* and the rare “short” isoform of the D2 receptor^12^. Due to this hybrid-like expression, we name it the D1H SPN.

Our results reveal an important organizing principle of neuronal identity and circuits. Continuous variation of identity has been observed previously to relate to spatial location in simple systems comprised of single cell types or single spatial axes^15–18^. Here we show that cell types are subdivided by multiple, independent spatial gradients of expression. We further show that the genes underlying these spatial axes can be conserved between discrete subtypes, allowing for the inference of spatial position of multiple cell types in complex tissues from a small number of reference markers. This also provides a fundamental explanation for why combinations of genes often are required to specify neuronal cell types: a cell is best described by the combination of its discrete identity and its position along each independent spatial axis. This discovery has obvious implications for the choice of strategy to establish comprehensive cell atlases of model organisms.

More broadly, our results have implications for neuronal circuits and computation. We find that neurons of the same type have highly localized gene expression patterns due to multiple spatial gradients of gene expression. We find that previously mapped connectivity correlates well to these gene expression patterns, suggesting that neuronal circuits can be subdivided into very small units depending on gene expression. This contrasts with existing models of neuronal circuits, which are traditionally defined by large groups of cells with shared regulatory regions from single marker genes. Our results suggest that developing combinatorial, spatially-precise approaches to targeting neurons may substantially improve our understanding of neuronal computation.

## Results

We previously found that most cells clearly classified as D1 or D2 SPNs^12^. However, we also reported several novel subtypes that co-expressed *Tac1* and *Penk* (D1-*Pcdh8*, D2-*Htr7*) as well as continuous gradients of expression within the major D1 and D2 subtypes. Importantly, we observed that some of subtypes seemed more discrete and defined by binary on/off patterns of gene expression, and other heterogeneity was more continuous and defined by gradients of gene expression. However, we were limited in our ability to clearly distinguish these distributions and determine the exact number of intermediates by our limited number of cells. We therefore prepared 1343 single dissociated D1-tdTom+ or D2-GFP+ SPNs from 5 adult mouse brains using Smart-Seq2 full-length amplification, and analyzed 1211 cells with at least 1e5 mapped reads and 1000 detected genes (Sup Fig 1A,B).

### Improved detection of discrete SPN types

We applied a standard clustering pipeline, PCA of the top variable genes followed by tSNE projection and Louvain clustering on a shared-nearest-neighbor (SNN) graph. This pipeline produced two apparently discrete clusters on a tSNE mapping. We first assessed the continuity of the clusters using partial approximate graph abstraction (PAGA)^19^. We initially found that none of the clusters separated into discrete types (Fig 1A-i). However, the gene expression underlying the D1/D2 difference appeared to change sharply (Fig 1B, Sup Fig 1E). Previously, we showed that the gene expression distribution underlying the D1/D2 separation was strongly bimodal and nearly all cells could be classified as either D1 or D2 ^12^. Additionally, we showed that we could considerably decrease the amount of technical variability in a PCA dimensionality reduction by truncating the PC loadings so that only the top PC-associated genes contributed to PCA (truncated PCA)^20^. We applied this modification to PCA and PAGA, and this resulted in a clear, discrete separation between D1 and D2 SPNs (Fig 1A-ii). We found evidence that this adjustment reduced the average contribution of batch (sequencing plate) variability to PCs (Sup Fig 1L), possibly because batch effects are caused by small changes consistent across thousands of genes. We also removed PCs that likely did not represent stable cell identity, like immediate early gene expression or batch effect (Sup Fig F-I). After this computational optimization, we identified 4 discrete clusters: D1 SPN, D2 SPN, a subtype expressing *Drd1a* and *Rreb1* that we identified as SPNs located in the islands of Calleja (ICj), and a subtype expressing *Nxph4* and *Pcdh8* that we had previously reported (D1-Pcdh8) (Fig 1B). We then re-applied dimensionality reduction and clustering to each of the 4 discrete subtypes. We found that truncated PCA-based clustering did not reveal any further discrete subtypes; however, truncated PCA of D2 subtypes considerably reduced batch effect (Sup Fig 1J). We compared cluster marker genes to a publicly-available RNA ISH database^40^ to map the heterogeneity within D1 and D2 SPNs to anatomical regions of the striatum (Fig 1C-F, Sup Fig 2A-E). Within D1 SPNs, we found 8 subgroups of cells, and within D2 SPNs, we found 7 subpopulations, nearly all of which corresponded to anatomical regions of the striatum (Fig 1C, D). We found a small number of cells expressing *Serpinb2* in D1 SPNs, which the *in situ* database showed to be highly specific to a novel sub-region of the medial shell (Fig 1C-G), Sup Fig 2A,E).

**Figure 1.**
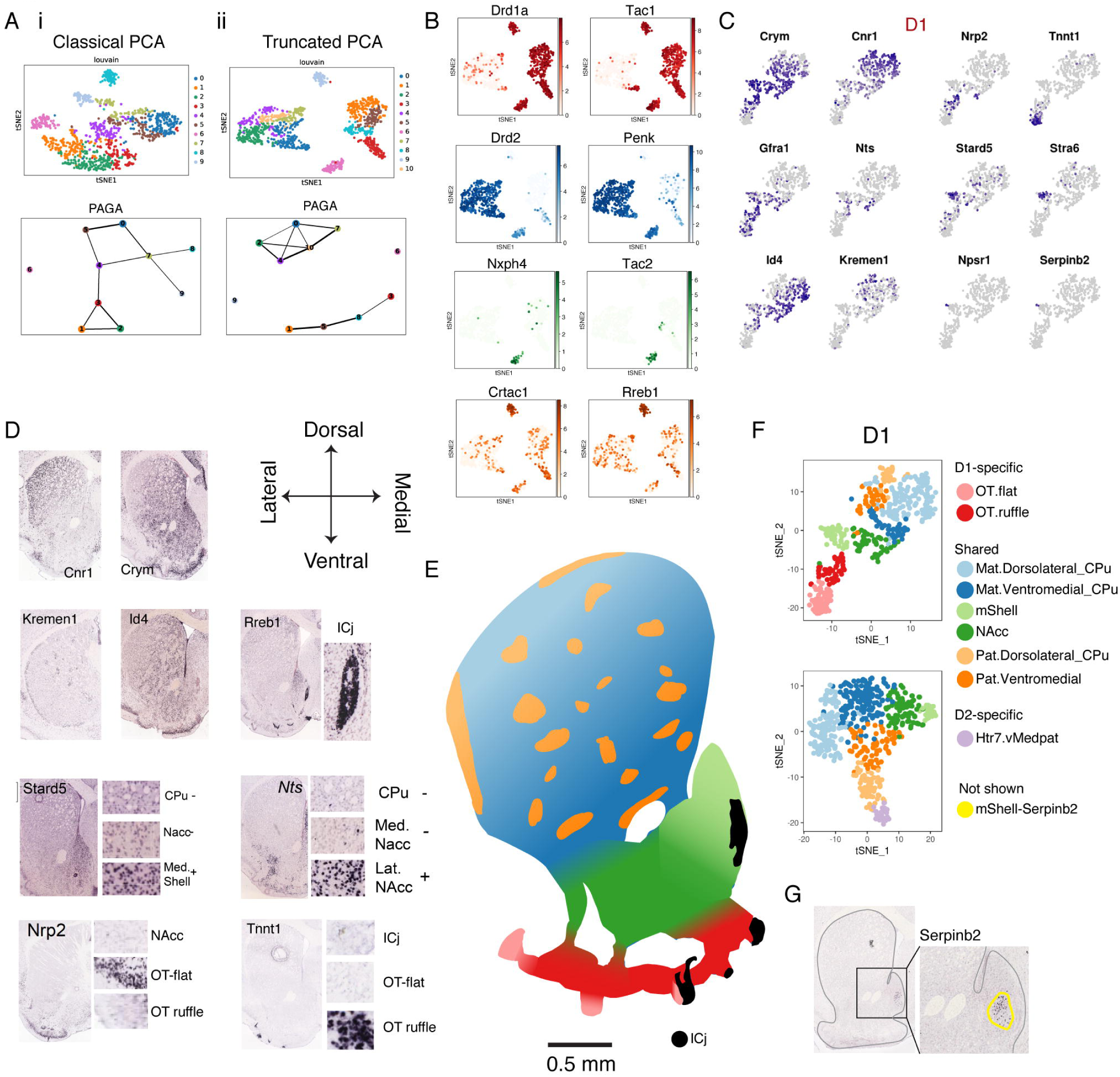
Computational analysis and mapping of SPN heterogeneity. A. SPNs clustered and discreteness analyzed with PAGA i) using regular PCA or ii) after truncating the PC loadings so that only 200 genes had non-zero loadings per PC. B. Gene expression defining 3 discrete subtypes: D1 (Drd1a and Tac1), D2 (Drd2 and Penk), and D1H (Nxph4 and Pcdh8). C. Subclustering D1 MSNs and markers defining each subcluster. Number of unique genes calculated as the number of genes enriched (FDR < 0.05) in only that subtype and no others. Several subtypes are clearly more “intermediate” and lack many clear unique markers. D. Mapping D1 and D2 subclusters to anatomical location using the Allen Brain Atlas *in situ* database. E. A map of the striatum with colors corresponding to the D1 subclusters (corresponding D2 map in Sup Fig X). F. Anatomical subclusters of D1 and D2 SPNs. Most subclusters are conserved, but several are unique to either D1 or D2. G. A novel subregion of the medial shell of the striatum marked by Serpinb2 expression (Allen Brain Atlas). scRNAseq plots are in Sup Fig X.

### Mapping the discrete cell types of the striatum

We then sought to map the discrete cell types of the striatum and understand whether their *in situ* distributions were also discrete. We quantified the discrete identity of SPNs by RNAscope *in situ* hybridization of their markers. We first stained and quantified D1 vs. D2 identity by the log ratio of *Tac1* to *Penk* expression (Fig 2A). As expected, D1 and D2 cells were spatially intermingled throughout most of the striatum (Fig 2B). We found that this identity was very strongly bimodal with few intermediates, corresponding to the discrete nature of D1 vs. D2 identity (with a small number of co-expressing cells that correspond to other subtypes) (Fig 2C).

**Figure 2.**
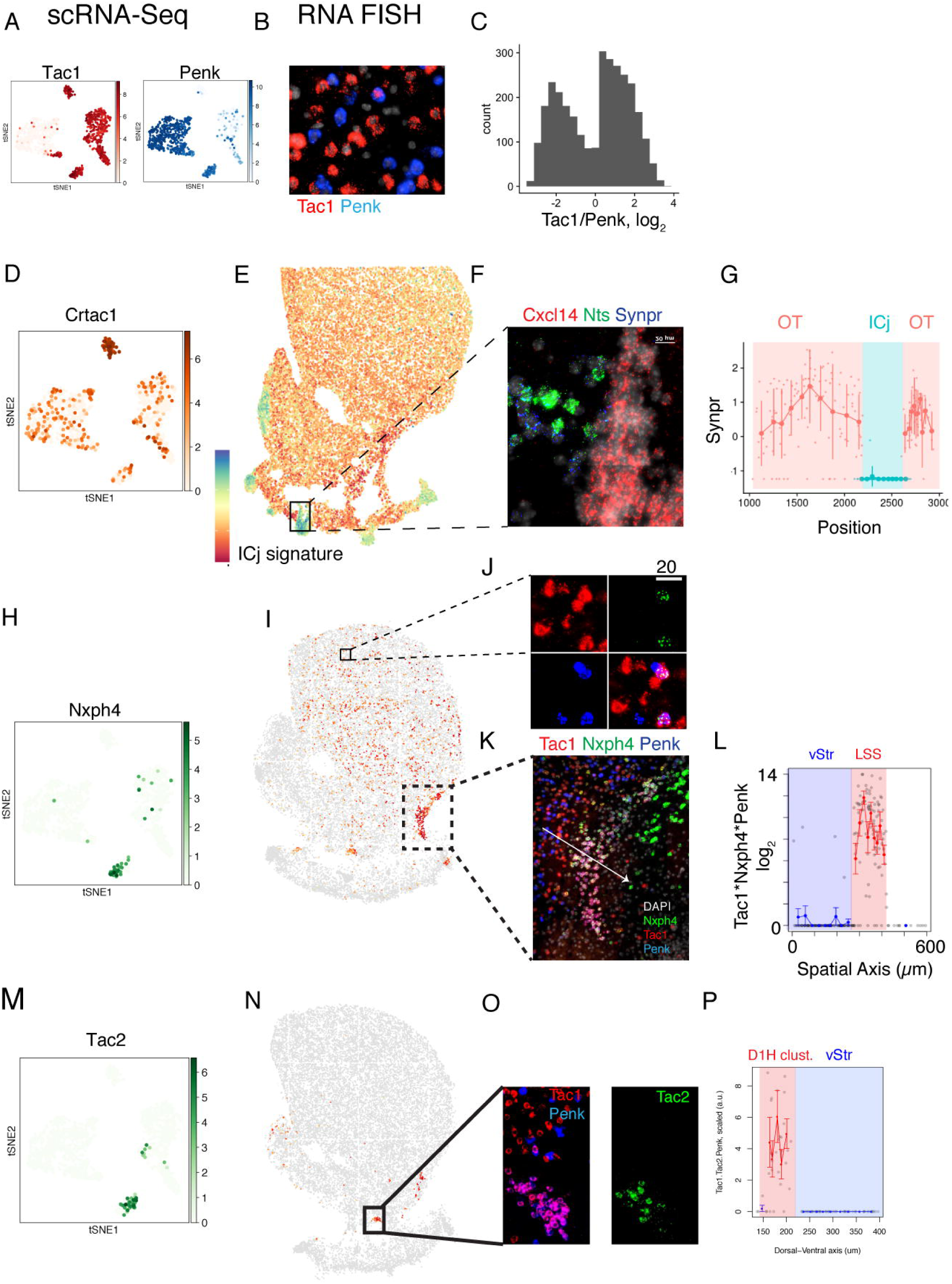
In situ mapping of the discrete types of SPNs. A. *Tac1* and *Penk* primarily separate discrete D1 and D2 SPNs. B. Fluorescence image the D1 and D2 discrete types using RNA in situ (RNAscope) of *Tac1* (red) and *Penk* (blue) mRNA. C. Distribution of *Tac1* vs. *Penk* expression, measured by the ratio of the integrated intensity per cell of either stain in log space. The distribution is strongly bimodal. D. Expression of *Crtac1*, which defines the discrete ICj subtype. E. *In situ* expression (RNAscope) of a signature corresponding to the ICj subtype. This signature is calculated as the ratio 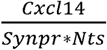, using integrated intensities per cell of each probe in log space. F. Fluorescence image of Cxcl14 (red), Nts (green), and Synpr (blue) in situ probes used to calculate the signature in 2F. Scale bar is 10µm. The change from D1-OT to ICj is step-like. G. *In situ* quantification of *Synpr* expression, showing a discrete, step-like change from the OT region to the ICj region. H. Expression of *Nxph4*, which defines the Nxph4+ subset of the discrete D1H subtype. I. *In situ* expression (RNAscope) of a signature corresponding to the D1H-Nxph4 subtype. This signature is calculated as *Nxph*4 * *Tac*1 * *Penk*, using integrated intensities per cell of each probe in log space. Any cell that does not express all three genes is colored grey. J. Fluorescence image of Tac1 (red), Nxph4 (green), and Penk (blue) in situ probes used to calculate the signature in 2F. Scale bar is 10µm. The D1H-Nxph4 subtype appears scattered throughout the CPu. K. Fluorescence image of Tac1 (red), Nxph4 (green), and Penk (blue) in situ probes in the lateral NAcc and lateral stripe of the striatum (LSS). Arrow corresponds to the spatial axis showing a discrete change in expression from the NAcc to the LSS. L. Single-cell quantification of the D1H-Nxph4 signature *Nxph*4 * *Tac*1 * *Penk*, in log intensity space, over the spatial axis defined by the arrow in Fig 2K. M. Expression of *Tac2*, which defines the Tac2+ subset of the discrete D1H subtype. N. *In situ* expression (RNAscope) of a signature corresponding to the D1H-Tac2 subtype. This signature is calculated as *Tac*2 * *Tac*1 * *Penk*, using integrated intensities per cell of each probe in log space. Any cell that does not express all three genes is colored grey. O. Fluorescence image of Tac1 (red), Tac2 (green), and Penk (blue) in situ probes in the NAcc and a ventral cluster. P. Single-cell quantification of the D1H-Tac2 signature *Tac*2 * *Tac*1 * *Penk*, in log intensity space, over the spatial axis defined by the arrow in Fig 2K.

We then analyzed the spatial distribution of the small discrete populations in our analysis. Our computational analysis predicted that the islands of Calleja (ICj) SPN subtype was discretely separated from the rest of the SPN types (Fig 1A-ii). Interestingly, this subtype is not present in previously published single-cell atlases of the nervous system. We quantified the identity of the ICj subtype as *Synpr - (Cxcl*14 + *Nts)* in log intensity space (Fig 2E). We found that expression of this identity changed in a discrete step along a spatial axis from the olfactory tubercle to the ICj (Fig 2F, G). This matches with previous anatomical observations that the islands of Calleja appeared distinct from the rest of the striatum, supporting the validity of our scRNAseq and *in situ* analyses^21^.

Finally, we stained for the Tac1/Penk-co-expressing D1H subtype, which is comprised of two continuous subtypes (*Nxph4*+ or *Tac2*+). D1H-*Nxph4* cells were intermingled with classical D1 and D2 cells in the dorsal striatum and clustered in ventrolateral structures (Fig 2I-K). We quantified D1H identity *in situ* as *Nxph*4 + *Tac*1 + *Penk* in log-intensity space and found a discrete step change in D1H identity from the ventral striatum to the D1H cluster (Fig. 2L). We performed a similar analysis for the Tac2+ D1H subtype, and saw it exclusively in discrete ventromedial clusters (Fig 2N-P). The D1H subtype had been previously mapped *in situ* but the discrete ventral clusters were not reported, likely due to the gene *Otof* not being a unique marker for D1H cells in the ventral striatum (Sup Fig 3C)^10^. Generally, we find that discrete gene expression patterns are associated with intermingled (D1, D2, D1H) or discretely-clustered (D1H, D1-ICj) neuronal types.

### Gene expression gradients underlie continuous subtypes of the striatum

We applied PC truncation, filtering, and PAGA to D1 and D2 SPNs to reveal the continuous axes of heterogeneity (Fig 3A, B). Generally, we found that continuous subtypes were spatially-adjacent. For example, the subtype mapping to the medial shell was continuous only with the NAcc subtype and not the CPu, which corresponds to the fact that the medial shell is physically adjacent only to the NAcc (Fig 3C, D). This led us to believe that the gene expression distributions underlying continuous subtypes represented gradients of expression. We found a signature of genes correlated to *Cnr1* and *Crym* shared between 3 discrete subtypes in the scRNAseq data (Fig 3E). *In situ* staining of Cnr1 and Crym revealed a gradient consisting of a 250-fold change in relative expression along the full 2mm-long dorsolateral-ventromedial axis (Fig 3F). Individual cells in between the dorsolateral-ventromedial poles had clearly intermediate expression levels of both *Cnr1* and *Crym* (Fig 3G, H). We describe the first single-cell *in situ* proof that these genes change along a dorsolateral-ventromedial gradient at a single-cell level and are not the result of heterogenous distributions of discrete cell types. Similarly, we found that the medial shell gene program was a gradient from the NAcc to the dorsal tip of the medial shell (Fig 1F, Fig 3C, Sup Fig 3D,E). The medial-shell subtype is not apparent in previous droplet-microfluidics single-cell atlases of the striatum, possibly due to dissociation bias.

**Figure 3.**
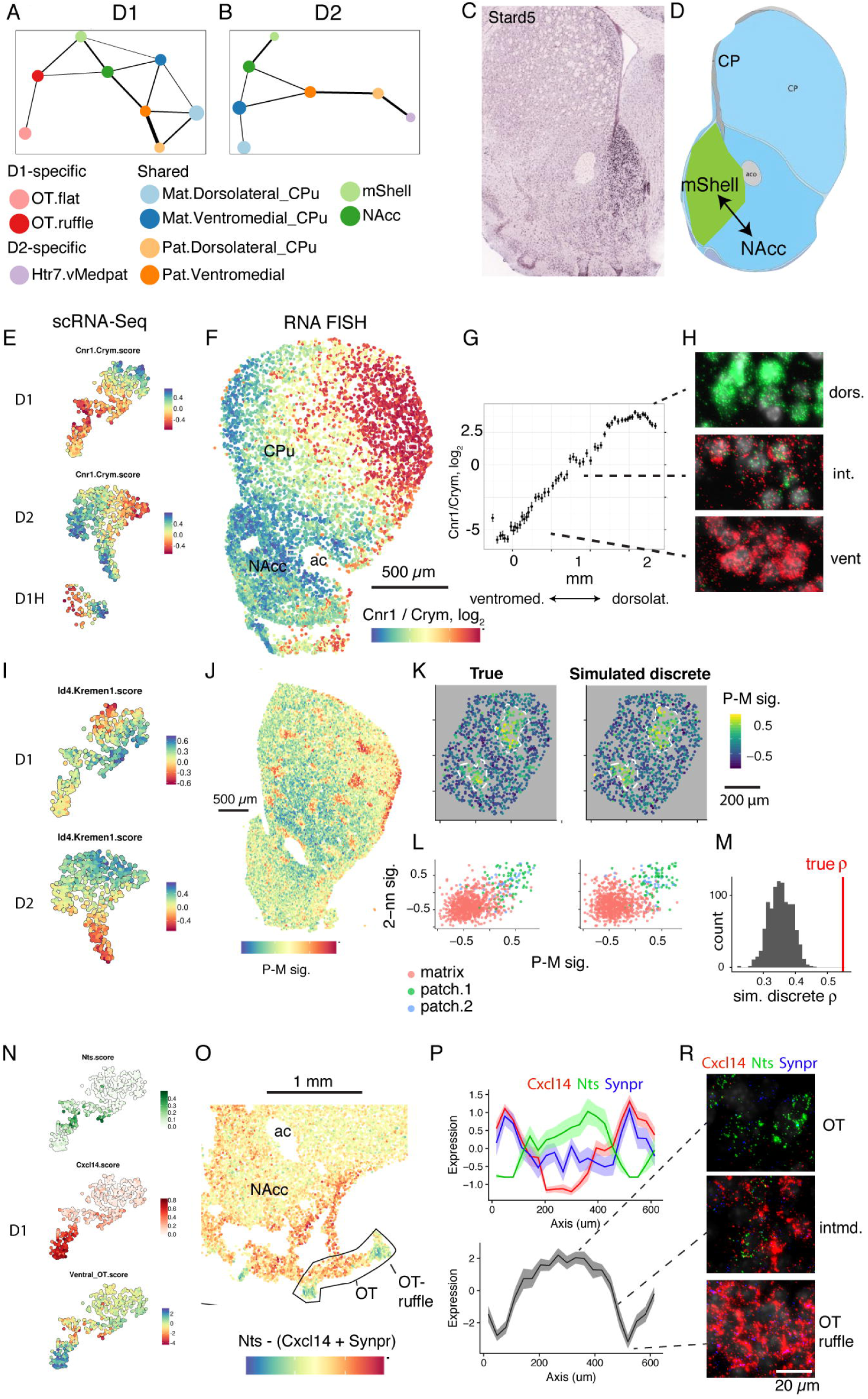
Independent spatial gene expression gradients underlie continuous subtypes of the striatum. A. PAGA of D1 SPNs, performed using 60-gene truncated PCA after filtering pheno-orthogoal PCs. B. PAGA of D2 SPNs, performed using 60-gene truncated PCA after filtering pheno-orthogoal PCs. C. Expression of the marker defining the medial shell (mShell) region of the striatum. D. Reference map of striatal anatomy showing that the mShell is anatomically adjacent to the NAcc, but not to the CPu E. Expression of a conserved *Cnr1-Crym* score across 3 discrete subtypes. The signatures were calculated by summing over the top 10 genes correlated to Cnr1 or Crym in all 3 subtypes. Each cell was scored the subtracting the *Crym* signature from the *Cnr1* signature. F. *In situ* quantification of *Cnr1* -*Crym*, using integrated probe intensity in log space. G. Quantification of the *Cnr1* - *Crym* along the dorsolateral-ventromedial axis of the CPu, defined by the white lines in Fig 3F. H. Representative in situ fluorescence of Cnr1 (green) and *Crym* (red) RNAscope probes in the dorsolateral (top), intermediate (middle), and ventromedial (bottom) CPu. I. Expression of a conserved *Kremen1-Id4* score across 2 discrete subtypes. The signatures were calculated by summing over the top 10 genes correlated to *Kremen1* or *Id4* in both subtypes. Each cell was scored the subtracting the *Id4* signature from the *Kremen1* signature. J. In situ quantification of the patch-matrix signature defined as (*Kremen1* + *Sema5b*) – *Id4*, using integrated probe fluorescence intensity in log space. K. Quantification of the patch-matrix signature within two real, adjacent patches (left panel). White lines show manually-defined boundaries of the patch, dividing the plot into 3 regions (patch.1, patch.2, and matrix). Right panel shows simulated discrete patches, where the expression of cells are scrambled within each of the 3 regions. L. Quantification of spatial information by the correlation of the patch-matrix signature of a cell to its 2^nd^ nearest-neighbor (2-nn). M. Monte Carlo simulation of the 2-nn correlation (rho) of 1000 discrete patches (grey distribution) and the true rho (red line). N. scRNAseq expression of *Nts* and *Cxcl14* signatures in D1 SPNs, calculated by the average value of the top 10 genes correlated to *Nts* and *Cxcl14*, respectively. Bottom panel represents the sum of the *Cxcl14* and *Synpr* signature subtracted from the *Nts* signature. O. *In situ* quantification of the OT-ruffle signature defined as *Nts* – (*Cxcl14* + *Synpr*), using integrated fluorescence intensity in log space. P. Quantification of individual genes (upper plot) and *Nts* – (*Cxcl14* + *Synpr*) (lower plot) along the spatial axis from a medial OT-ruffle to the flat OT to a lateral OT-ruffle. Q. Representative in situ fluorescence of *Cxcl14* (red), *Synpr* (green), and *Nts* (blue) RNAscope probes in the flat OT (upper), intermediate (middle), and OT-ruffle (bottom).

We then found that genes correlated to the patch-matrix markers *Kremen1* and *Id4* formed a gradient in both D1 and D2 cells that was largely orthogonal to the *Cnr1*-*Crym* gradient (Fig 1D, Fig 3I). We stained for the patch-matrix organization using probes to *Kremen1, Sema5b* (which was highly correlated to *Kremen1*), and *Id4* (Fig 3I). We quantified patch-matrix identity as *Kremen*1 + *Sema*5*b - Id*4 in log-intensity space. The patches appeared to lack discrete boundaries, but the 2D spatial arrangement was too complex to easily map to a single spatial axis (Fig 3J, K). Therefore, to determine whether the patch-matrix organization was discrete or not, we manually identified boundaries of the patches and performed a Monte Carlo simulation of discrete patches (Fig 3K). We quantified the degree of spatial organization as the correlation between a cell and its second-nearest-neighbor (*ρ*_2*nn*_) (Fig 3L). We found that the patch-matrix was much more spatially organized than a discrete-patch model would predict (*p* < 0.001) (Fig 3L, M). This indicated that patch-matrix identity is not binary and instead depends continuously on distance from the center of a patch. We thus present the first proof that the patch-matrix distribution is a continuous spatial relationship at the single cell level.

We also observed continuous subtypes that predicted anatomical location over very small length scales. One of the D1-only continuous subtypes localized to the “ruffles”, small protrusions of the olfactory tubercle. In the scRNA-seq data, a small subgroup of cells expressing *Nts* was most directly continuous with the D1 population expressing *Cxcl14*, a marker enriched in the OT ruffles (Fig 1D, Fig 3N). To determine if these were gradients, we stained for *Cxcl14, Nts*, and *Synpr* (a gene de-enriched in the OT) (Fig 3O, P). We found that *in situ* expression levels of these genes changed continuously from the ventral NAcc to the OT (Sup Fig 3F, G). Their expression again changed continuously along a spatial axis from the flat OT to the OT ruffles over just 150 µm (Fig 3G, H). This demonstrates that continuous relationships in scRNAseq data predict anatomical position even over very small length scales. We also identified a novel expression gradient within the OT-ruffles where *Tnnt1* increased with proximity to the pial surface only in the lateral OT-ruffles (Fig 1D). Previous single-cell atlases of the brain did not analyze the ventrolateral and OT regions of the striatum^10^ and therefore omitted these subtypes.

### A novel spatial compartment of the striatum: D2H ventromedial “patches”

As reported previously, a subtype of D2 SPNs (D2-*Htr7* or D2H) co-expresses *Penk* and *Tac1*^10,12^. Unlike the D1H population, this population exclusively expresses *Drd2* and not *Drd1a*. Our scRNAseq analysis predicts that it is continuous with the D2-patch cells, but not the D2-matrix cells (Fig 4A). This is supported by the fact that there is strong overlap between marker genes defining the D2-patch and D2H cell types (Fig 4A). Since the D2-patch cells have a distinct spatial distribution, and we have shown that continuous subtypes are always anatomically related, we hypothesized that the spatial distribution of the D2H cells would also be patch-like. We thus stained for *Th, Htr7*, and *Tac1*. *Htr7*+/*Tac1*+ cells appeared in patchy distributions, but more ventromedial than most of the patches marked by *Kremen1* or *Sema5b* (Fig 3J, Fig 4B, C). *Htr7*+/*Tac1*+ cells were 14x more likely to be *Th*+ than *Htr7*-/*Tac1*+ cells, matching the scRNA-Seq data (Fig 4A,D). Our scRNAseq analysis predicted that *Th* would mark only the most distinct tip of the D2H continuum (Fig 4A). Indeed, triple-positive cells were limited to the most ventromedial part of the D2H-patches, particularly the medial shell (Fig 4B). We therefore define a novel spatial compartment of the striatum composed of ventromedial patch-regions containing *Tac1*+/*Htr7*+ D2 SPNs, and a sub-region expressing *Th* that is restricted to patches in the medial shell. This subtype of SPNs has been observed in our and other single-cell databases, but its anatomy and relationship to the striatal patch compartment has not been described^10^. Intriguingly, the *Th*+ cells did not express *Ddc*, suggesting that they do not metabolize dopamine but instead use tyrosine hydroxylase in a different pathway.

**Figure 4.**
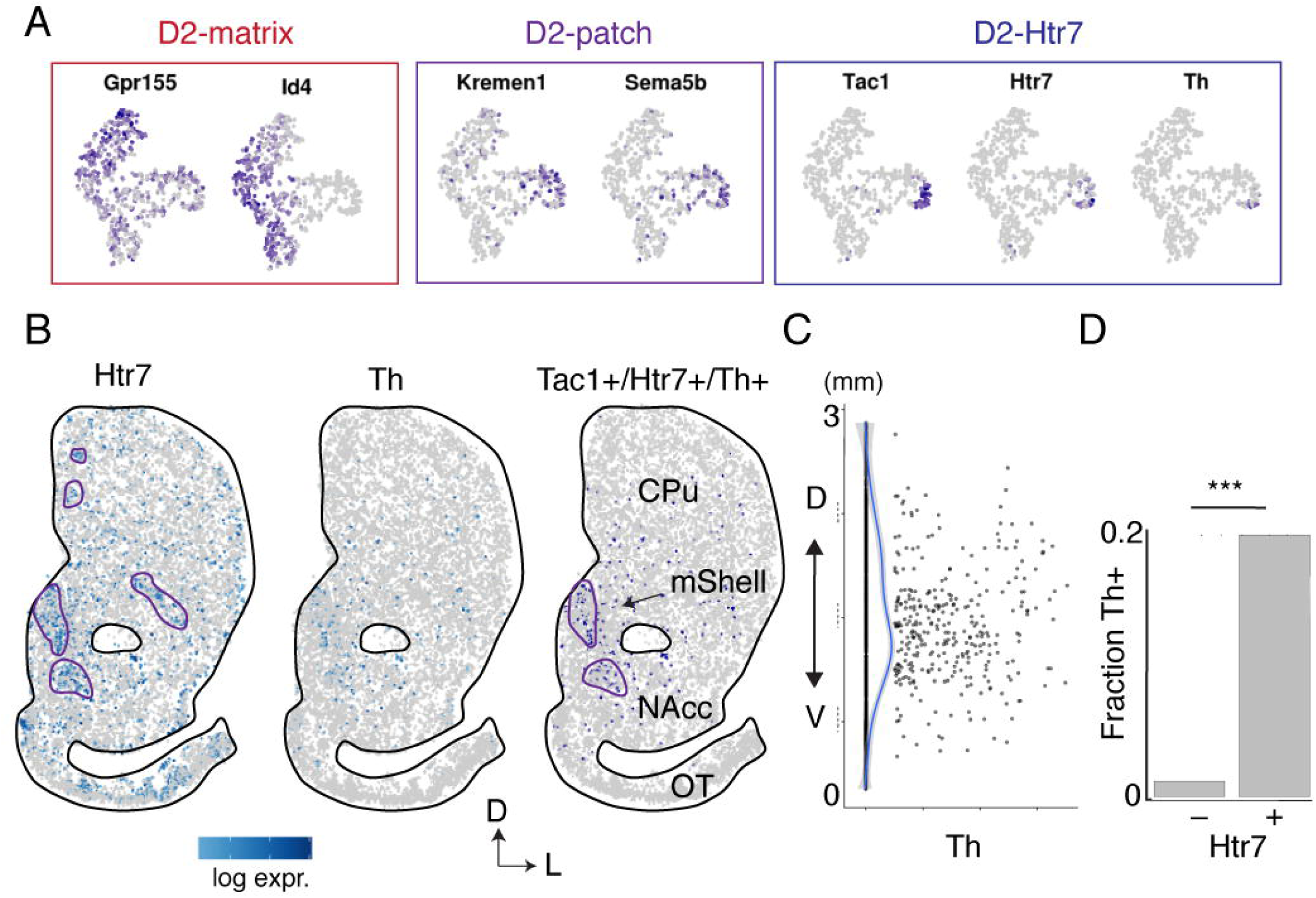
Defining a novel region of the striatum populated by patch-like D2H SPNs. A. ScRNAseq expression of matrix, patch, and D2H markers in D2 SPNs (tSNE dimensionality reduction). The D2H subtype is seen as a continuous extension of the D2-patch system. B. *In situ* quantification of D2H markers Htr7 (left), Th (middle). Cells below the detection threshold are grey. Triple-positive cells are shown in the right plot displayed as blue and non-triple-positive cells are grey. C. Quantification of Th expression among Tac1+ cells showing a strong enrichment in the middle of the striatum, corresponding to the location of the medial shell. D. *In situ* quantification of the fraction of cells expressing Th, separated by whether they also express Htr7. P < 0.001, OR = 14.

### Splicing and identity determination of a D1R/D2R co-expressing type

Though D1 and D2 receptors are primarily non-overlapping in expression and function, there is known to be co-expression and even heterodimerization of these two receptors (Frederick et al., 2015). We found that most D2 SPNs expressed the D1 receptor at scattered, low levels and the D1 SPNs expressed almost no D2R (Fig 1C). The subtype with highest co-expression of *Drd1a* and *Drd2* was the D1H-*Nxph4* subtype (Fig 1C). Drd1a/Drd2 co-expression in this subtype was not observed in recent droplet microfluidics atlases, possibly due to the low detection rate of those technologies^10,22^. The D2 receptor is expressed in two isoforms, short (D2S) and long (D2L)^23^ (Fig 5B). We found that just 8% of isoform-specific reads mapped to D2S in D2 SPNs, consistent with previous literature (Fig 5C)^23^. Expression was similar in the novel ventromedial-patch subtype D2-Htr7, which co-expresses *Tac1* and *Penk*, but not Drd1a and Drd2 (Fig 1B, Fig 5C). Intriguingly, we found that 34% of isoform-specific reads mapped to D2S in the D1H-*Nxph4* subtype (Fig 5C). Therefore we find that the D2S isoform is specific to D1R/D2R co-expression.

**Figure 5.**
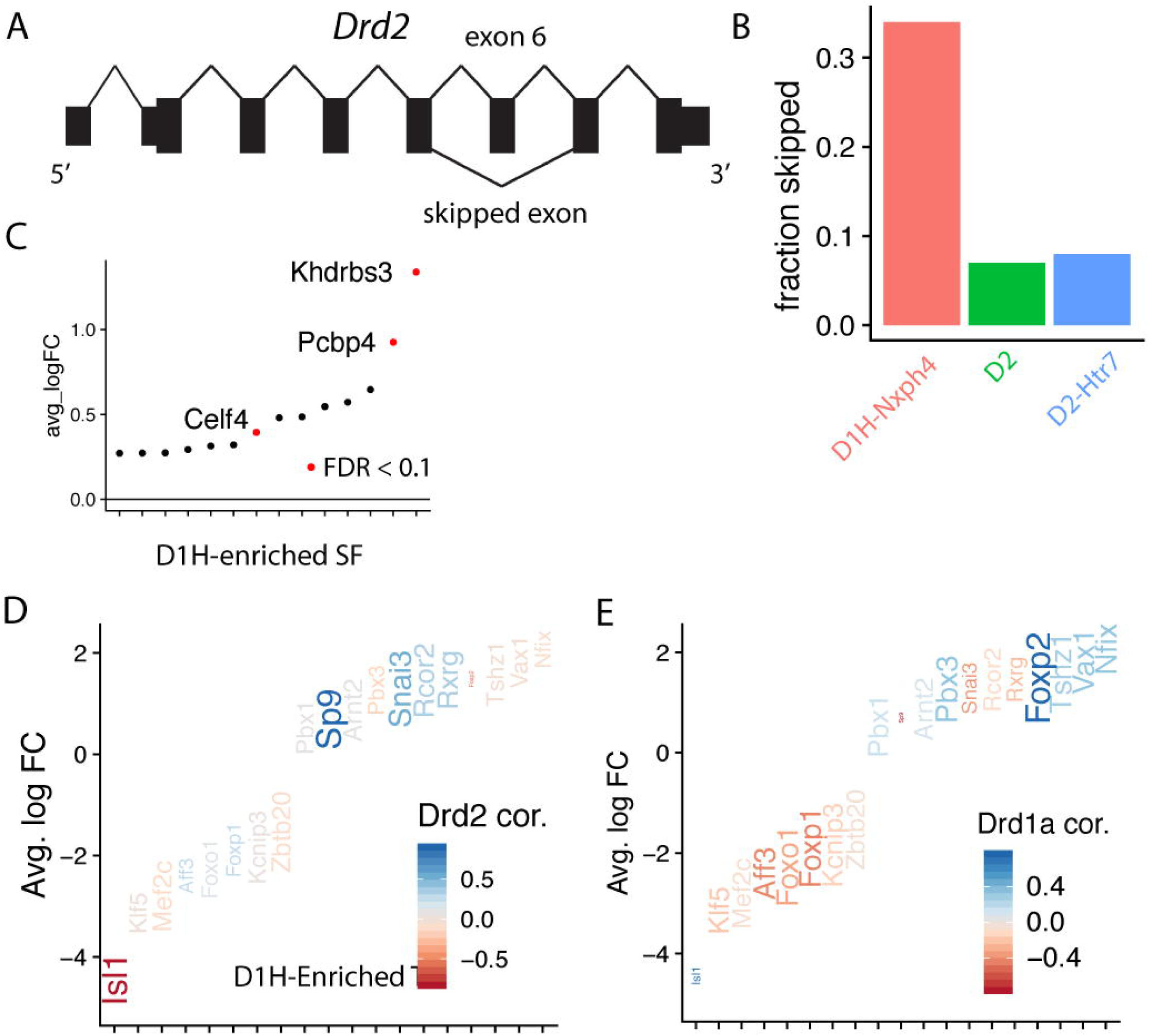
Splicing and identity determination of a D1R/D2R co-expressing type of SPN. A. Genomic structure of the D2 receptor. The 6^th^ exon is known to be skipped. B. Quantification of the fraction of reads aligning to the skipped exon over the total reads aligning to either junction, 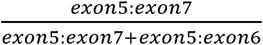. C. Average log fold change of top differentially expressed splice factors (selected from GO: 190641), comparing D1H-Nxph4 cells to all the other SPNs. The 3 genes significantly enriched in D1H-Nxph4 are displayed as text and in red (FDR < 0.1). D. Transcription factors (TFs) that are candidates for controlling *Drd2* expression. TFs shown are all those significantly enriched in D1H-Nxph4 cells vs all other SPNs (FDR < 0.1). Genes are colored by their correlation to *Drd2* () across all SPN subtypes. Gene size is calculated as 0.5 + *Drd2* correlation * sign(D1H-Nxph4 log FC). Genes that are positively correlated to *Drd2* and enriched in D1H-Nxph4 are large; genes that are negatively correlated to *Drd2* and de-enriched in D1H-Nxph4 are large; others are small. E. Transcription factors (TFs) that are candidates for controlling *Drd1a* expression. TFs shown are all those significantly enriched in D1H-Nxph4 cells vs all other SPNs (FDR < 0.1). Genes are colored by their correlation to *Drd1a* across all SPN subtypes. Gene size is calculated as 0.5 + *Drd1a* correlation * sign(D1H-Nxph4 log FC). Genes that are positively correlated to *Drd1a* and enriched in D1H-Nxph4 are large; genes that are negatively correlated to *Drd1a* and de-enriched in D1H-Nxph4 are large; others are small.

To understand the molecular pathways driving this phenomenon, we investigated transcription factors (TFs) and splicing factors (SFs) enriched in the D1H-*Nxph4* subtype. We found 3 SFs enriched in D1H-*Nxph4* (FDR < 0.1), one of which may be responsible for splicing out exon 6 (Fig 5C). We found 19 TFs enriched (FDR < 0.1) (Fig 5D). To determine which TFs might control Drd2 expression in D1H-*Nxph4*, we also measured the correlation of each TF to *Drd2* across all SPNs. *Isl1* was the most highly anticorrelated to *Drd2* throughout all SPNs and de-enriched in the D1H-*Nxph4* subtype. *Sp9* was the most highly correlated to *Drd2* and enriched in the D1H-*Nxph4* cell type. This suggests that *Isl1* is an inhibitor and *Sp9* is an activator of *Drd2* expression in D1H-*Nxph4* cells. *Sp9* and *Isl1* are required for proper formation of D2 and D1 neurons^24–26^. However, since *Isl1* is not expressed in D1H-Nxph4 cells, it cannot be solely responsible for turning on *Drd1a* expression. Therefore we measured the correlation of each enriched TF to *Drd1a*, and found that *Foxp2* was by far the most positively-correlated to *Drd1a* among TFs enriched in D1H-*Nxph4* cells. Together, our results suggest *Isl1* as a *Drd2* repressor, *Sp9* as a *Drd2* activator, and *Foxp2* as a *Drd1a* activator.

## Discussion

Striatal projection neurons have previously been classified into D1 and D2 SPNs with regional differences defined by anatomical region. Recently, two additional types of SPNs have been found and partially characterized *in situ*^10,12^. We demonstrate that these identities are composed of four discrete types, D1, D1H, D2, and ICj neurons. Within the striatum, D1, D2, and D1H SPNs are composed of continuous subtypes that correspond to anatomical gradients and compartments.

The effort to describe neuron types is still lacking an ontology that can describe the full range of neuronal heterogeneity while providing structure and clarity. In particular, it remains difficult to relate high-dimensional molecular heterogeneity to well-mapped anatomical regions. Early efforts to use single-cell sequencing classified neurons into many discrete types, and mapped the specific regions in which those clusters were present^22,27^. However, it was observed that many neurons had characteristics of at least two classes^11^. It was also observed that similar or overlapping neuron classes were often anatomically adjacent^11,22,28^. Additionally, it has been found that neurons from adjacent regions are often transcriptionally similar^23^. *In situ* staining showed that gene expression could vary in a gradient-like manner across an anatomical region^17^. In simple systems, this meant that dissociated cells could be mapped at high resolution to their position along a spatial axis just by their transcriptome^2,7,16,29^. However, these results have not been combined into a broad framework that can be applied to more complex systems.

Our work addresses this challenge by providing a clear framework for relating neuronal identity and anatomy, and for revealing in greater detail than ever before how discrete and continuous heterogeneity relates to anatomy. In the striatum, we showed that discrete types have no implied spatial relationships to one another, and can coexist randomly intermingled throughout a region as in the case of D1 or D2 neurons, or can exist in discrete regions composed of single subtypes, as in the Islands of Calleja. Stable continuous heterogeneity almost always encodes spatial gradients. Subtypes that are directly continuous are generally anatomically adjacent. Previous papers have reported similar subtypes that are anatomically adjacent, but that appear discrete due to a small number of co-expressing neurons^30^. We instead propose that spatial gradients can occur over very small length scales, exemplified by the OT-ruffle transition, which occurs over just 150 µm. Additionally, continuous heterogeneity does not always imply an obvious linear gradient of expression, as demonstrated by the patch-matrix axis. Our conclusion is that stable, continuous neuronal subtypes are related by an underlying anatomical axis, regardless of the details of the continuity; discrete subtypes are not.

Our work provides a natural explanation and solution to a key challenge of how to define neuronal subtypes. We show that each neuron is defined by multiple independent axes of heterogeneity, which demonstrates that neurons are best defined by a combinatorial identity. The effort of assigning neurons to types has been hindered by the difficulty of finding unique marker genes for each subtype^28,31^. Previous papers have dealt with this by naming neuron types by the combination of genes that best define them ^10^, although this can lack clarity and interpretability. We showed that both the genes encoding dorsoventral gradients and the genes encoding the patch-matrix gradients are shared between the D1 and D2 discrete subtypes. Because of this, there are not great marker genes defining, for example, just the cells of the dorsal D1 patches. Instead, these cells are best defined by three genes: a D1 gene (*Drd1a*), a dorsal gene (*Cnr1*), and a patch gene (*Kremen1*). Therefore we conclude that most projection neurons in the striatum in fact are best defined by three values: one discrete value indicating D1 or D2 identity, one continuous value indicating dorsoventral identity, and one continuous value indicating patch-matrix identity. Our work therefore describes both an interpretable ontology, and a fundamental biological reason for using it.

The dorsolateral-ventromedial gradients of expression align with the topology of striatal connectivity, which also changes in a graded manner along the dorsolateral-ventromedial axis^32,33^. In fact, most of the anatomical compartments we describe have distinct patterns of connectivity, and connectivity changes in a graded manner along the dorsolateral-ventromedial axis^32,33^. This suggests that circuits are themselves not discrete units composed of large numbers of cells. Instead, circuits may be defined highly locally, by the combinatorial and gradient identities of their underlying neurons.

Our *in situ* characterization of the newly-discovered D1H subtype reveals that it has a complex anatomical distribution. Surprisingly, we find that it is present in homogenous clusters of cells in the NAcc, and comprises the entirety of the lateral stripe of the striatum. As observed previously, it is also scattered throughout the CPu^10^. The scattered dorsal D1H subtypes expresses the unique marker *Nxph4* as well as the conserved dorsal marker *Cnr1*, whereas the D1H subtype in the ventral clusters expresses the unique marker *Tac2* and the conserved ventral marker *Crym*. The ventral clusters and LSS largely correspond to the TAC2+ clusters found by immunohistochemistry^34^. The D1H cells also have a third subtype expressing the conserved medial shell marker *Stard5*; we therefore predict that this represents D1H cells present in the medial shell region.

Interestingly, we show that the D1H subtype co-expresses the D1 and D2 dopamine receptors. We also show that this co-expressing type is the only one with high levels of the D2S isoform of the D2 receptor. Previous studies have shown that the D2S isoform only acts presynaptically^23,35^. Separate studies have proposed the existence of a D1/D2 heterodimer and suggested that it is only localized somatically and presynaptically^36^. Since D1/D2 co-expression and D2S isoform expression are correlated, both transcriptionally and physiologically, the D1/D2 heterodimer may require the D2S isoform to form. The difficulty of assessing D1/D2 co-expression or heterodimerization could thus be due to the alternate D2 isoforms having lower binding to common D2 antibodies.

We utilized the complex expression of Drd1a and Drd2 in our dataset to discover their transcriptional regulators. *Isl1* and *Sp9* were highly correlated to D1/D2 identity, as previously reported^12^, and *Isl1* is known to be required for SPN development. However, both of them explained *Drd2* expression much better than *Drd1a* expression, suggesting that they are more likely regulators of Drd2. Drd1a expression was much better explained by *Foxp2*. *Foxp2* has been shown to control *Drd1* expression in the songbird striatopallidal system, and the gene is among the most highly conserved among mammals^37,38^. Interestingly, it is the only gene clearly linked to acquisition of language in humans^39^. Our studies provide motivation for studying the role of *Foxp2* in the learning and the dopamine receptive system.

Combining in situ and single-cell data, we discovered a novel anatomical compartment of the striatum. The D2 subtype expressing *Htr7* and co-expressing *Tac1* and *Penk* was recently described in single-cell data^10,12^. Interestingly, we find that this subtype is continuous with the D2-patch cells in the scRNA-seq data. We then showed that it was present in patchy regions in the ventromedial CPu and medial shell. We thus conclude that this subtype is a ventromedial extension of the patch-matrix complex, and define the *Drd2*+ ventromedial patch as a novel region of the striatum. The patch region of the striatum was previously only observed in the CPu and NAcc and not in the medial shell^40^. Our results suggest that this is because the D2-*Htr7* subtype has a different gene expression signature than the D1- and D2-patch types.

Overall, our work advances our understanding of the basis of neuronal heterogeneity and provides many targets for investigating the role of the striatum in health and disease. Our work has broad implications for studies of neural circuits and neural computations. Clearly the impressive local heterogeneity of neurons of a given classical type such as D1 SPNs means that even manipulations of small numbers of neurons of a class will in fact affect sets of different neurons, and will have different significance based on where exactly that manipulation is applied. Traditionally circuits are explored by probing hundreds, often thousands of neurons that are thought to be the same type, and correlating their activity with a downstream readout. However, if neurons have a defined identity that goes beyond their type and is determined by location and gene expression patterns, it is difficult to envision defined circuits between large numbers of cells. There must be a substructure to these circuits that depends on the precise identity of the neurons involved, and this substructure may be as important for neural computations as the overall connectivity of the neurons themselves. It is possible that a redefinition of the concept of neural circuits may lead to a better understanding of how the brain processes information by analyzing the brain’s computational networks not as bulk circuits, but deconvolving synaptic connections into more precise point-to-point circuits that differ among neurons of the same type depending on their location and gene expression patterns.

