## Supplementary Methods and Figures for "Discrete and Continuous Cell Identities of the Adult Murine Striatum"

#### Animals

Generation of D1-tdTom/D2-GFP, D1-tomato, and Aldhl1-GFP BAC reporter mice was previously described [1,2]. All lines were maintained by backcrossing to C57Bl/6J. All other experiments were conducted on 5-7 week old male C57/Bl6 mice from Jackson Labs. Mice were weaned at ~21 days of age and housed in groups of 2-5 on a 12-hour light/dark cycle (lights on 0700-1900h), with free access to food. All procedures conformed to the National Institutes of Health Guidelines for the Care and Use of Laboratory Animals and were approved by the Stanford University Administrative Panel on Laboratory Animal Care.

#### Tissue Dissociation

Parasagittal slices (350 µm) containing striatum were prepared using standard procedures. Briefly, mice were anesthetized with isoflurane and decapitated, brains were quickly removed and placed in ice-cold low-sodium/high-sucrose dissecting solution. Slices were cut by adhering the lateral surface of the brain to the stage of a Leica vibroslicer, and placed in ice-cold hibernate-A solution (Gibco, Themo Fisher). The striatum was dissected from each slice, and the tissue was dissociated using papain-based dissociation. Briefly, striatal tissue was incubated with frequent agitation at 30°C for 20 min 2mg/ml papain solution in Hibernate-A, followed by washing with hibernate-A including 1mg/ml Ovomucoid protease inhibitor (Sigma). The striatal slices were triturated briefly using fire polished glass Pasteur pipettes until a single-cell suspension was obtained. The suspension was cleaned from small debris by centrifuging for 3 min at 450g through a three-step density gradient of Percoll (Sigma), top layer containing small debris was discarded and remaining cells were pelleted for 5 min at 550g. The pellet was resuspended in 0.1 ml of hibernate A with DAPI (4',6-diamidino-2-phenylindole, 1:4000 dilution; Sigma) to label dead cells[5]. Striatal tissue was kept in ice-cold solution throughout the processing until loading onto the microfluidic chip, except during the 20 min dissociation with papain at 30°C.

#### Purification of Striatal Cells by FACS.

The dissociated cell suspension from adult D1-tdTom/D2-GFP or Aldhl1-GFP mouse was passed through a 70-µm cell strainer (BD Biosciences). Viable cells (DAPI negative) were sorted by FACS (Aria II; BD Biosciences). For specific enrichment of genetically labeled cell types, fluorescent striatal cells were analyzed to determine optimal parameters and gating for FACS. Finally, test sorted cells were imaged on confocal microscope as described previously[5]. Cells were sorted into ice-cold hibernate-A and pelleted by centrifugation for 5 min at 300g. Single cells were sorted into 96 well plates filled with 4 µl lysis buffer composed of 0.05% Triton X-100 (Sigma) and, ERCC spike-ins (1:24000000 dilution), 2.5 µM oligo-dT, 2.5 mM dNTP and 2U/µl of recombinant RNase inhibitor (Clontech) then spun down and frozen at -80°C. Plates were thawed and libraries prepared as described below.

#### Smart-Seq2 library preparation

96-well plates with sorted single cells were incubated for 3min at 72°C and immediately placed on ice. To perform reverse transcription (RT) we added each well a mix of 0.59µl $H_2O$, 0.5µl SMARTScribe™ Reverse Transcriptase (Clontech), 2µl 5x First Strand buffer, 0.25µl

Recombinant RNase Inhibitor (Clontech), 2µl Betaine (5M Sigma), 0.5µl DTT (100mM) 0.06µl MgCL2 (1M Sigma), 0.1µl Template-switching oligos (TSO) (100µM AAGCAGTGGTATCAACGCAGAGTACrGrG+G). Next reaction RT reaction mixes were incubated at 42°C for 90 min followed by 70°C for 5 min. Pre amplification of cDNA done by adding 12.5µl KAPA HiFi Hotstart 2x (KAPA Biosystems), 2.138µl $H_2O$, 0.25µl ISPCR primers (10 µM, 5′-AAGCAGTGGTATCAACGCAGAGT-3), 0.1125µl Lambda Exonuclease under the following conditions: 37°C for 30 min, 95°C for 3 min, 21 cycles of (98°C for 20 sec, 67°C for 15 sec, 72°C for 4 min), final extension at 72°C for 5 min. Cells were then cleaned using 2 rounds of AMPure bead (Beckman-Coulter) cleanup at a 0.8:1 ratio of beads to PCR product. The cleaned cDNA was then tagmented using the Nextera XT DNA kit and following the included instructions, using 11 cycles of PCR to amplify the tagmented cDNA (Illumina). Tagmented cDNA was then pooled and analyzed with a Bioanalyzer 2100 High Sensitivity DNA kit (Agilent) to assure that >80% of the product was in the 200-1000bp range. This process was repeated separately for each animal. Libaries were sequenced using 2x75 reads base pairs (bp) paired-end on Illumina HiSeq 2000 or NextSeq to a target depth of $1x10^6$ reads/sample.

**Processing and Quality Control of Single-cell RNA-seq Data**

Raw reads were aligned using rnaSTAR to the mm10 genome with *Tdtomato* and *Gfp* transgenic mRNA sequences and ERCC synthetic RNA added. Gene counts were determined using htseq-count with "strict" mode on (all ambiguous alignments discarded)[6] . Gene expression was quantified as the number of counts per gene divided by the total number of mm10 counts, respectively, times $10^6$ to get expression in counts per million (cpm). Values of log(cpm + 1) were used for analysis. Low-quality cells, which we defined as having < 1000 genes detected and $<10^5$ counts as mapped by HTSeq-count, were removed for downstream analysis.

*Normalization*

Cells were normalized by dividing by the total number of counts per cell, multiplying by $10^6$, and log-transforming with a pseudocount of one to make log cpm using NormalizeData in Seurat. Genes were scaled for PCA using the ScaleData function.

*Contaminant removal*

A first round of clustering was performed using default Seurat parameters for FindVariableGenes and RunPCA for dimensionality reduction on cells passing quality thresholds. A group of 4 cells with low PC9 scores were enriched for genes like *Apoe*, *Slc1a3*. Querying the brain atlas database mousebrain.org showed that these genes were highly specific to astrocytes, so these cells were determined to be likely contamination and removed from downstream analysis (Sup Fig 1C). Performing Seurat and PCA on the remaining cells revealed 3 cells expressing *Sp8, Nfib*, and other genes (Sup Fig 1D). Querying the single-cell atlas dropviz.org showed these to be neurogenic genes. These cells were removed from further analysis. No other PCs or cells were observed to be enriched for glial markers.

**Computational analysis of discrete cell types in scRNA-Seq data**

*Optimization of clustering of discrete types*

The remaining cells were clustered and visualized using the following Seurat-based pipeline to optimize PCA for cluster discreteness. PCA was run on variable genes chosen with FindVariableGenes default parameters. The JackStraw function was used to compute the significance of each PC; there was a large fall-off of significance after PC15, so the first 15 PCs were used for clustering and graph abstraction. Truncated PCs were computed by setting all gene loadings below the top $N$ positive and top $N$ negative loadings to 0; in Fig. 1 $N$=200. The top 15 default or truncated PCs were used as input in the shared nearest-neighbor (SNN) graph generation and tSNE visualization with RunTSNE, and Louvain clustering with FindClusters of the Seurat package version 2.3.4 and R version 3.5.1. The first 15 PCs generated by Seurat were saved and used to calculate the partial abstract graph approximation (PAGA) algorithm with the scanpy package[7] version 1.3.7 and Python version 3.6.6. The following sequence of commands using the PC scores as input: sc.pp.neighbors with n_neighbors=5, sc.tl.draw.graph with default parameters, sc.tl.louvain with resolution=0.5, sc.pl.paga with groups = 'louvain', and sc.pl.paga with threshold = 0.09.

*Filtering PCs that do not represent stable cell identity*

PCs were filtered by overlaying the PC scores on the tSNE plots (Sup Fig 1F), and overlaying the genes with top PC loadings on tSNE plots for each PC (Sup Fig 1G, H). Several PCs had high loadings of immediate early genes (IEG) like *Fos* and *Arc*, and those were removed because they are known to be highly transient[8] (Sup Fig 1F, H, I). Other PCs were dominated by very-high-expressed genes like *Lars2*, a long non-coding RNA, which had almost zero dropout and did not have expression patterns that correlated to any clear subtype separation (Sup Fig 1G).

*Determining discreteness with PAGA*

Discrete cell clusters were determined using the above pipeline, with the scanpy paramaters as described above, with truncated PCA with $N$=200 and after filtering out PCs that did not represent stable cell identity. We called cell types discrete if the PAGA-calculated edges connecting them were below 0.12. We used this threshold because manual inspection of PCs of cell types with a connecting edge width of ≤ 0.12 appeared discrete, whereas as cells with connecting edges > 0.12 were completely continuous (Sup Fig 1J,K).

**Mapping heterogeneity within D1 and D2 SPNs**

*Subclustering D1 and D2 SPN discrete types*

D1 and D2 SPNs were initially clustered using Seurat. Each discrete type was separately re-normalized and scaled with NormalizeData with scale.factor = 1e6 and ScaleData with default parameters. Highly variable genes filtered with FindVariableGenes with y.cutoff=.3. Any variable genes found to be expressed in >95% of cells was removed before PCA. PCA was calculated with RunPCA with default parameters. PCs were truncated as described previously, with $N=60$. As before, PCs not likely to correspond to stable identities or corresponding to one or two outlier cells were removed.

*Mapping subclusters to in situ expression atlases*

The subclusters of each discrete type were assigned based on the genes expressed in them. Cluster-defining genes were manually searched in the Allen Adult Mouse Brain Atlas (mouse.brain-map.org). Genes that were consistent between D1 and D2 SPNs and strongly-detected in the striatum were used to map subtypes (Fig 1C, D, Sup Fig 2A-E). Multiple genes per cluster or anatomical regions were referenced to ensure that the cluster assignments were valid.

*Generation of shared gene expression signatures*

A shared *Cnr1*-*Crym* expression signature was generated in the following way. The Spearman correlation of every gene to *Cnr1* and *Crym* was calculated separately within each of the D1, D2, and D1H SPN subtypes. Each gene was scored for *Cnr1* correlation by the sum of its 3 within-subtype correlations. The top 10 genes were chosen by this metric. The top 10 *Crym*-correlated genes were similarly chosen. The *Cnr1-Crym* score was computed as the sum of the top 10 *Cnr1* genes less the sum of the top 10 *Crym* genes within each cell. The genes used were the same across all 3 discrete subtypes, by design. The shared *Kremen1-Id4* signature was generated in the same way for the D1 and D2 SPNs. The *Nts* and *Cxcl14* scores were generated in the same way in D1 SPNs; this was not a shared signature as the olfactory tubercle cells were only readily detected in the D1 SPNs.

*Quantification of Drd2 splicing*

Bam files from 30 randomly-selected cells each from the D1, D2, and D1H SPN types were visualized with the Integrated Genome Viewer (IGV) software version 2.4.10 after loading the mm10 genome[9]. *Drd2* splicing was quantified using a Sashimi plot generated by IGV. The fraction of skipped reads was calculated as the ratio of reads mapping to the exon5-exon7 junction over the sum of the reads mapping to the exon5-exon6 and exon5-exon7 junction.

### Quantitative *in situ* mapping of SPN gene expression

*Generating fresh-frozen brain slices*

Wildtype C57B/L6 background mice were sacrificed at 8-12 weeks and perfused with ice-cold saline. The brain was removed and placed into a cryomold (VWR cat. no. 15160-215) filled with OCT (Tissue-Tek ®, VWR). This was then placed into an isopentane bath that was pre-cooled in liquid nitrogen. Care was taken to avoid splashing of any isopentane onto the exposed OCT surface. When the OCT was almost completely frozen (small drop of unfrozen OCT at the exposed surface) it was transferred to an insulated container filled with powdered dry ice, and stored at -80C for up to 3 months. The night before slicing, the brain was placed in a -20C freezer and allowed to equilibrate to -20C. The day of slicing, it was placed in a pre-cooled cryostat (Leica CM 3050 S) with the settings object temperature (OT): -12 to -15C; chamber temperature (CT) -25C. All chambers and tools were cleaned with RNAse-away (ThermoFisher cat. no. 10328011) and 70% ethanol (Sigma). A new blade was installed in the blade holder. The glass anti-roll plate was kept cold by placing dry ice wrapped in foil on top of it every ~5 slices. Slices were cut at 12-14 µm thickness with a quick, even movement of the blade. Brains were sliced coronally. Slices were flattened with a brush on the cold cutting surface, and mounted onto a Superfrost glass slide by bringing the slide into contact with the slice and allowing it to melt onto the slide, slowly, to avoid bubble formation. Each slice was quickly placed under a dissection microscope (Nikon) under dark-field illumination, and a photo was taken for reference (especially noting edge integrity and location of any bubbles). Slices that contained both the CPu and NAcc were kept for hybridization. Slides were stored at -80C for up to 6 months.

*RNAScope in situ hybridization*

All tools were cleaned with RNAse-away before staining. RNAScope probes (ACDBio) were hybridized following the manufacturer's instructions for fresh-frozen tissue. Briefly, brain slices were fixed in 4% PFA for 15 minutes. Then they were dehydrated in increasing concentrations of ethanol for 5 minutes each (70%, 85%, 100%, 100%). Then they were incubated for 30 minutes with Protease IV (ACDBio). Then they were hybridized to the target probes for 2 hours in an RNAScope hybridization oven at 40C (ACDBio). Then they were washed and incubated with the amplification reagents (Amp-1, Amp-2, Amp-3, and Amp-4) as per manufacturer instructions. They were then stained with DAPI (ACDBio) for 30 seconds and mounted with Prolong ™ Gold antifade reagent (ThermoFisher).

*Imaging*

Slides were imaged with a Zeiss AxioImager 2 using a 20X 0.8NA air objective (Zeiss). Multiple overlapping tile images were acquired. 10-12 z-stacks with 0.6-1µm spacing were used to assure that every part of the tile was in focus. Exposure times were 250-700ms. ZEN software (Zeiss) was used to do maximum intensity projection and stitching of the tiles using DAPI as the reference channel.

*Image analysis*

Images were analyzed with custom CellProfiler scripts[9]. Nuclei were smoothed with an 8-pixel artifact diameter with the Smooth function. 60-pixel speckle enhancement was done to reduce background, and then 8-pixel feature suppression to reduce the chance of segmenting on nucleoli. Nuclei were segmented with IdentifyPrimaryObjects with a typical pixel diameter between 35-80 pixels. Cells were segmented by expanding the nuclei segmentations by 7 pixels' radius with hole filling. Probe intensity per cell was then quantified with the following pipeline. Background and illumination variation was reduced on all probe channels with EnhanceSpeckles with a typical diameter of 30 pixels (roughly the size of a cell). Probe images were then manually thresholded to remove background. MeasureObjectIntensity was used to quantify probe intensity per segmented cell (Sup Fig 3A-B). Further analysis was done in custom R scripts. Gene expression was quantified as the log of the integrated intensity, with a lower cutoff manually determined by comparing to images to determine the level at which ~0-1 probes were found.

### Supplementary Figure 1

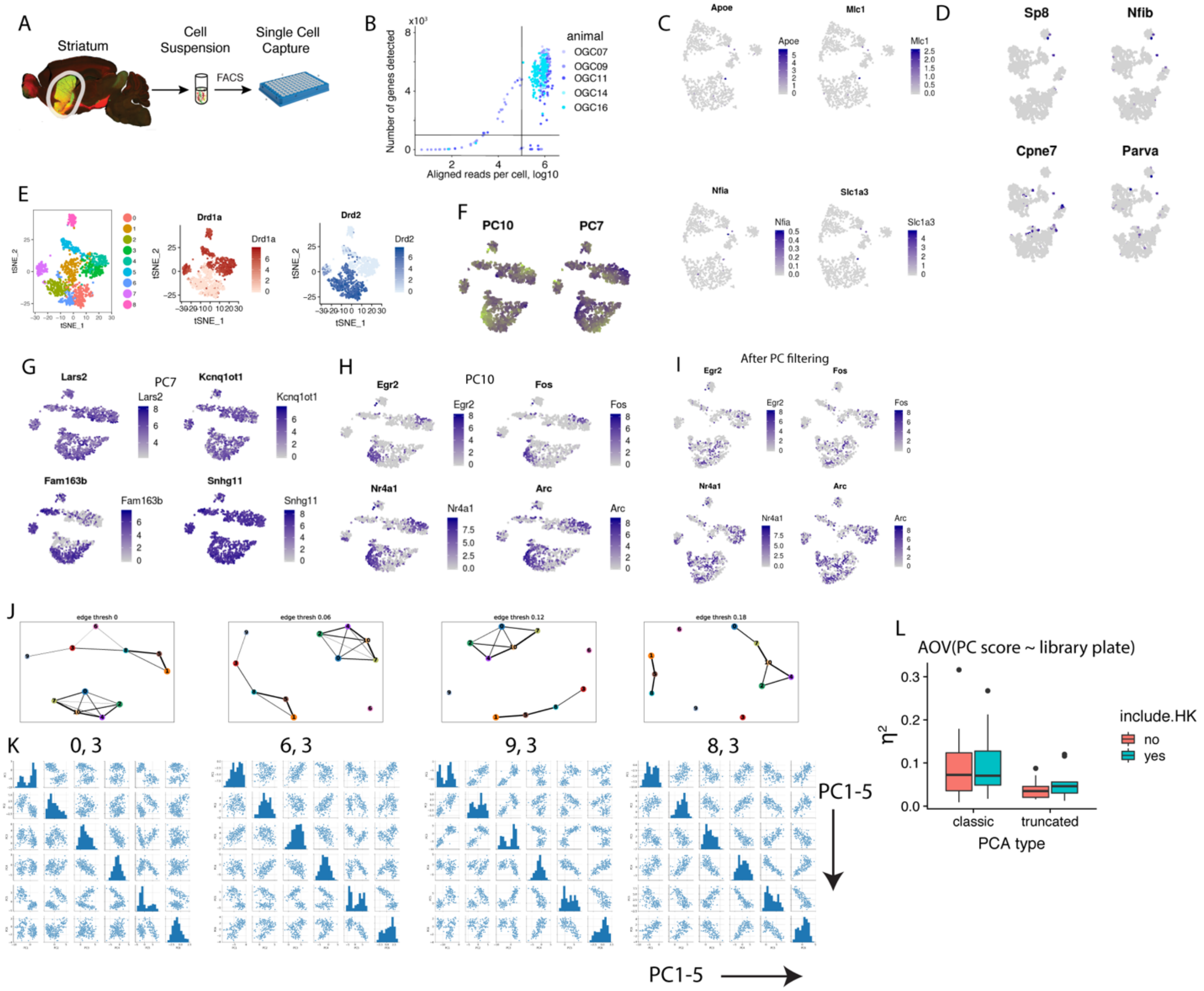


**Supplementary Figure 1: Improved pipeline for determining cell type continuity in single cell sequencing data**

A. Experimental outline for sorting D1+ and D2+ SPNs into 96-well plates for SmartSeq-2.

B. QC metrics and cutoffs for single-cell sequencing data.

C. Identification of astrocyte-contaminated cells.

D. Identification of neurogenic cells.

E. Classical PCA and tSNE of D1 and D2 SPNs overlaid with markers of discrete SPN subtypes (*Drd1a* and *Drd2*).

F. Overlaying truncated PC scores of PCs closely associated with IEGs (PC10) and housekeeping genes (PC7).

G. Top 4 genes by PC7 loading generated by PCA of all SPNs.

H. Top 4 genes by PC10 loading generated by PCA of all SPNs.

I. Immediate-early genes overlaid of tSNE of all SPNs after PC filtering and dimensionality reduction.

J. PAGA of SPNs with different values for edge thresholds. Clusters are generated by scanpy.

K. The PCs underlying an example pair of clusters whose edge is broken at the edge threshold in the PAGA image in (J).

L. The contribution of batch effect to the top 10 PCs measured by analysis of variance. Y-axis is $\eta^2$. Include.HK denotes whether high-expressed genes (expressed in >95% of cells) were included in the list of highly-variable genes included in PCA. PCA was run using Seurat RunPCA with genes chosen by FindVariableGenes with y.cutoff=0.5 on all D2 SPNs. The relationship is highly significant ($p=0.004$, $n = 20$). The decrease in batch effect $\eta^2$ is positively correlated to the original batch effect size (Pearson correlation = 0.96).

**Supplementary Figure 2**

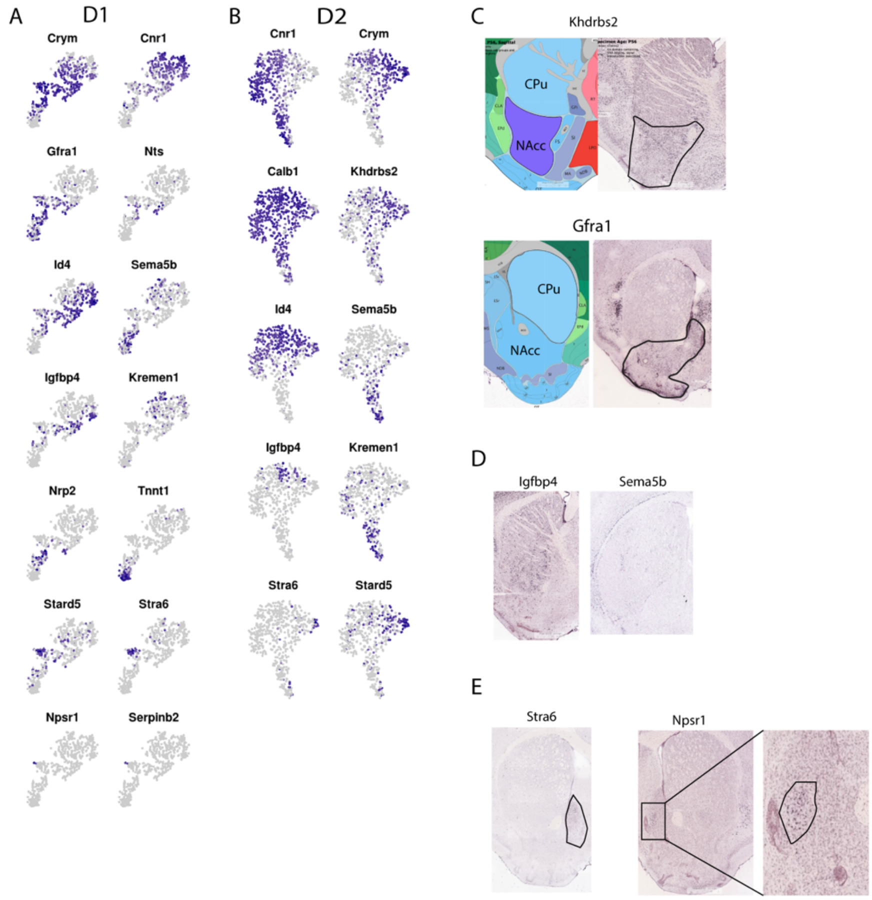


**Supplementary Figure 2. Assignment of spatial subtypes of SPNs using marker genes.**
A. Spatial marker genes overlaid on tSNE of D1 SPNs.
B. Spatial marker genes overlaid on tSNE of D2 SPNs.
C. *In situ* expression in the Allen Mouse Brain Atlas of marker genes used to define the NAcc SPNs in D1 (*Gfra1*) and D2 (*Khdrbs2*) SPNs.
D. *In situ* expression of additional marker genes used to define patch and matrix SPNs that were not included in Figure 1.
E. *In situ* expression of additional marker genes used to define medial shell and novel *Serpinb2*+ medial shell subregion SPNs that were not included in Figure 1.

**Supplementary Figure 3**

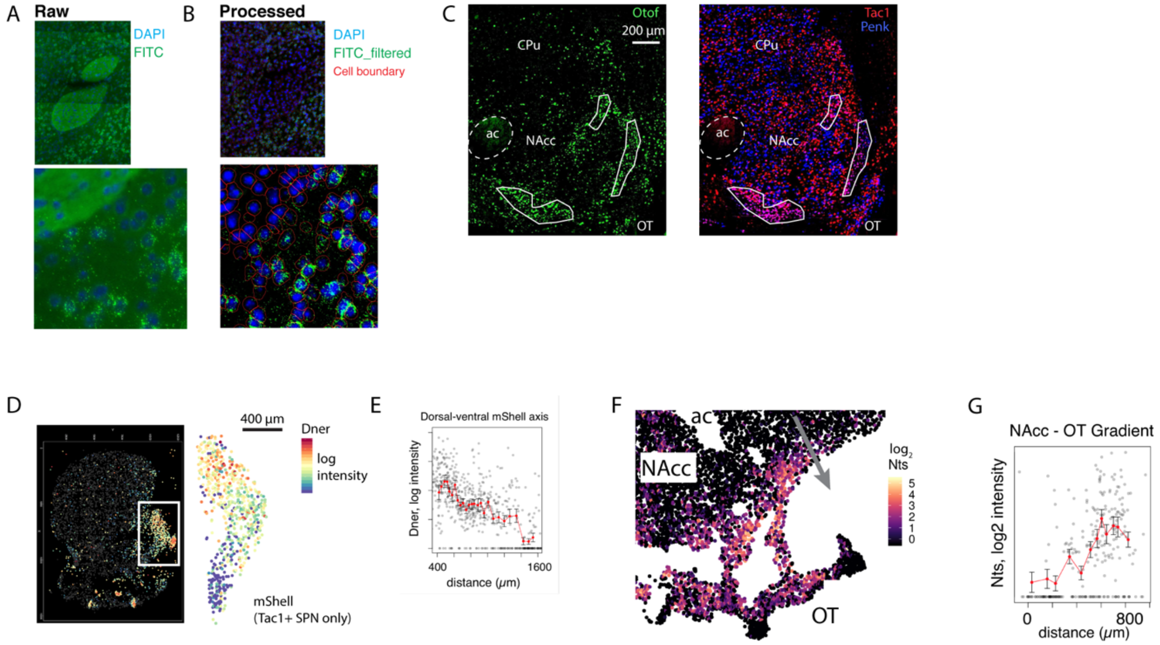


**Supplementary Figure 3. Quantification of gene expression *in situ*.**
A. Example raw image of an RNAscope positive control probe in the FITC channel and DAPI in the blue channel in a fresh-frozen adult mouse brain tissue slice.
B. Example processed image of *in situ* staining with an RNAscope control probe showing computational elimination of background and reduction of illumination variation in the FITC probe channel, and cell boundary generation in red using the DAPI channel to segment nuclei.
C. Expression of *Otof*, a previously-described marker of D1H cells, alongside *Penk* and *Tac1*, in the ventral adult mouse striatum.
D. Expression of *Dner* in the medial shell of the mouse striatum; expression is log integrated probe intensity.
E. Quantification of *Dner* expression along the ventral-dorsal axis within the medial shell.
F. Expression of *Nts* in the transition region between the NAcc and olfactory tubercle.
G. Quantification of *Nts* expression along the axis defined by the grey arrow in (G), in the transition region between the NAcc and the olfactory tubercle. See Sup Figure 2A for the scRNA-seq expression pattern of *Nts.*
